# From Orphan Genes to Cryptogenic Gene Candidates: Reassessing Uniqueness

**DOI:** 10.1101/2025.06.28.662130

**Authors:** Sheetalpreet K. Maan, Xuan Y. Butzin, Steven X. Ge, Nicholas C. Butzin

## Abstract

**Background:** Orphan genes lack recognizable homologues outside a given taxonomic unit; thus, they have elusive evolutionary origins. They have long been invoked to explain lineage specific traits in medicine and evolutionary biology. Here, we re-analyzed a 2023 repository of putative orphan genes compiled from over 80,000 bacterial species. Using iterative homology-based analyses, we reassessed each gene’s taxonomic distribution across a broader genomic landscape.

**Results:** These results challenge the long-held assertion that species truly harbor large numbers of orphan genes and instead demonstrate that their prevalence has been overestimated. Many “orphan genes” from 2023 now match homologues in other bacterial taxa (2025), demonstrating that sparse sampling had inflated earlier orphan counts. To better reflect these findings, we propose the term Cryptogenic Gene Candidates (CGCs). This offers a more precise alternative to orphan genes, as it more accurately reflects the provisional and potentially temporary nature of these genes’ apparent uniqueness. This revised terminology acknowledges that many genes identified as having a unique origin may lose this status as more genomes are sequenced. Although our analyses substantially reduced the number of false-positive CGCs, it cannot determine whether a given CGC truly encodes a functional protein or is an artifact of bioinformatic analysis. To minimize annotation artifacts, we then applied additional computational filters to prioritize candidates most likely to encode bona fide proteins.

**Conclusions:** This work revises the understanding of orphan gene prevalence, establishes CGCs as a more accurate classification, and provides a prioritized set of candidates to guide future experimental studies.

## Introduction

Most genes have homologs in other organisms. However, orphan genes (also called ORFans or Taxonomically Restricted Genes (TRGs)) have been defined as genes with no detectable homology outside a specific species or lineage^1^. These genes are believed to contribute to speciation and evolution as they enable organisms to undergo adaptations, helping them to survive environmental changes^2^. There are several mechanisms reported in the literature that explain the origin of such genes, including horizontal gene transfer, duplication and divergence, and *de novo* origination^3-5^. Interestingly, orphan genes are observed across various life forms, such as vertebrates^6,7^, insects^8^, nematodes^9^, yeast^10^, plants^11^, and bacteria. Unfortunately, due to the absence of evolutionary history, most orphan genes remain functionally unannotated, making it difficult to understand their biological significance. Hence, many predicted orphan genes have hypothetical status. This knowledge gap prompted us to investigate the roles of orphan genes, aiming to infer their potential functions. Also, hypothetical genes are a part of “dark genome,” which consists of the poorly understood, unannotated, or previously overlooked regions of an organism’s genome that lack clear functional annotations. Orphan genes remain an intriguing area of study because they constitute a substantial portion of the dark genome. Recent research has revitalized efforts to determine whether the hypothetical genes are functional and, if so, what traits they encode. A study showed that dark genes have high mutation rates in certain cancers and impact patient survival^12^. Their critical vulnerabilities suggest promising therapeutic targets, emphasizing the need for further research.

In this study, we investigated orphan genes across 10 different bacterial species and demonstrated that as more genomes are sequenced, previously identified orphan genes lose their unique status. This pattern indicates that orphan genes are frequently overestimated, likely due to the limited and biased sample of genomes currently available. Given the results from this work, we prefer to use the term *Cryptogenic Gene Candidates* (CGCs) (a term we justify in the manuscript), as many genes previously classified as orphan genes are not truly unique to a lineage. We anticipate that with continued sequencing efforts, more genes will similarly lose their orphan status.

Building on these findings, we introduce a methodology leveraging computational tools to prioritize CGCs for experimental investigation to infer their roles. This offers a framework to understand these enigmatic genetic elements better as sequencing technologies continue to advance.

## Results and Discussion

### Advanced Screening Approach Led to a Reduction in the Number of Potential Orphan Genes

Orphan genes differ from non-orphan genes in their taxonomic distribution. While non-orphan genes can be found across multiple species, genera, or other taxonomic units; orphan genes are typically confined to a specific taxonomic unit. If an orphan gene is restricted to a single genus, it is classified as a genus-level orphan gene, whereas if it is found only within a particular species, it is categorized as a species-level orphan gene (**Fig. 1a**). In this study, we focused on species-level orphan genes from selected bacterial species. These genes were distributed across various genomic locations, underscoring the complexity associated with their presence and organization (**Fig. 1b**). To distinguish genuine orphan genes from potential artifacts of bioinformatic predictions, we reassessed the orphan genes reported in TRGdb^13^ using the BLASTp^14^ approach.

**Fig. 1.**
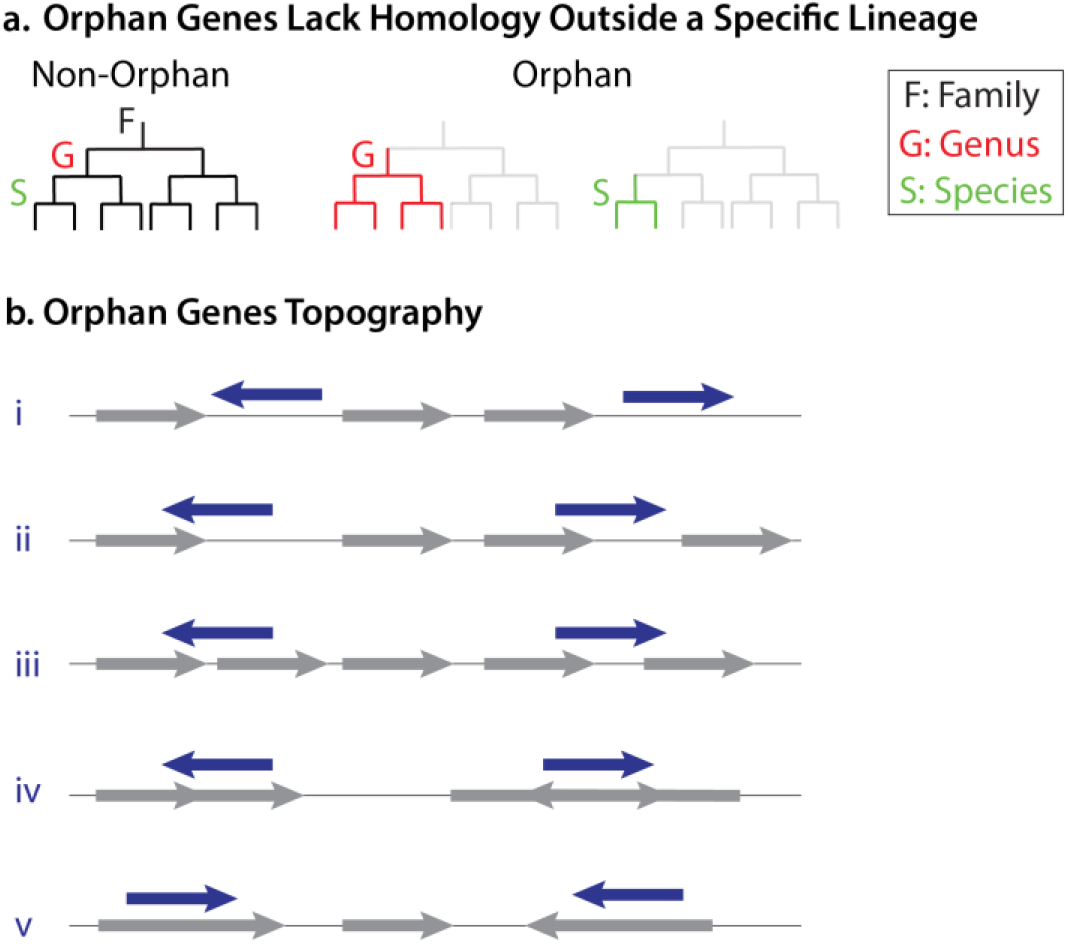
Classification and Genomic Topography of Orphan Genes. **a**. Distinguishing orphan gene from non-orphan genes. Orphan genes are characterized by their restricted taxonomic distribution. If a gene is present at the genus level but absent outside that genus, it is classified as a genus-specific orphan gene. Similarly, a species-specific orphan gene is identified only within a single species and not beyond. These genes exhibit no detectable homology outside their respective taxonomic levels. **b**. Genomic topography of orphan genes. Orphan genes can be found in various genomic locations: **i**. intergenic regions (between two genes), **ii**. overlapping a single gene, **iii**. overlapping two genes, **iv**. overlapping two overlapping genes, and **v**. located within a gene. Some of the genes were already annotated in the NCBI data while others (the majority) were unannotated.

The analysis covered 10 bacterial species: *Escherichia coli, Salmonella enterica, Klebsiella pneumoniae, Pseudomonas aeruginosa, Enterococcus faecalis, Bacillus subtilis, Mycobacterium tuberculosis, Helicobacter pylori, Legionella pneumophila*, and *Chlamydia trachomatis*. Our results demonstrated that most of the orphan genes lost their status in the case of all bacterial species due to the detection of significant similarity in other bacterial genera, families, or taxonomic units. Consequently, it led to their exclusion from further study. Also, the BLASTp approach employed 4 steps (E value: 1E-3) to ensure the reliability of our results (**Fig. 2a**). Each step was designed to progressively refine our identification of true orphan genes by checking for sequence similarity across different levels of relatedness.

**Fig. 2.**
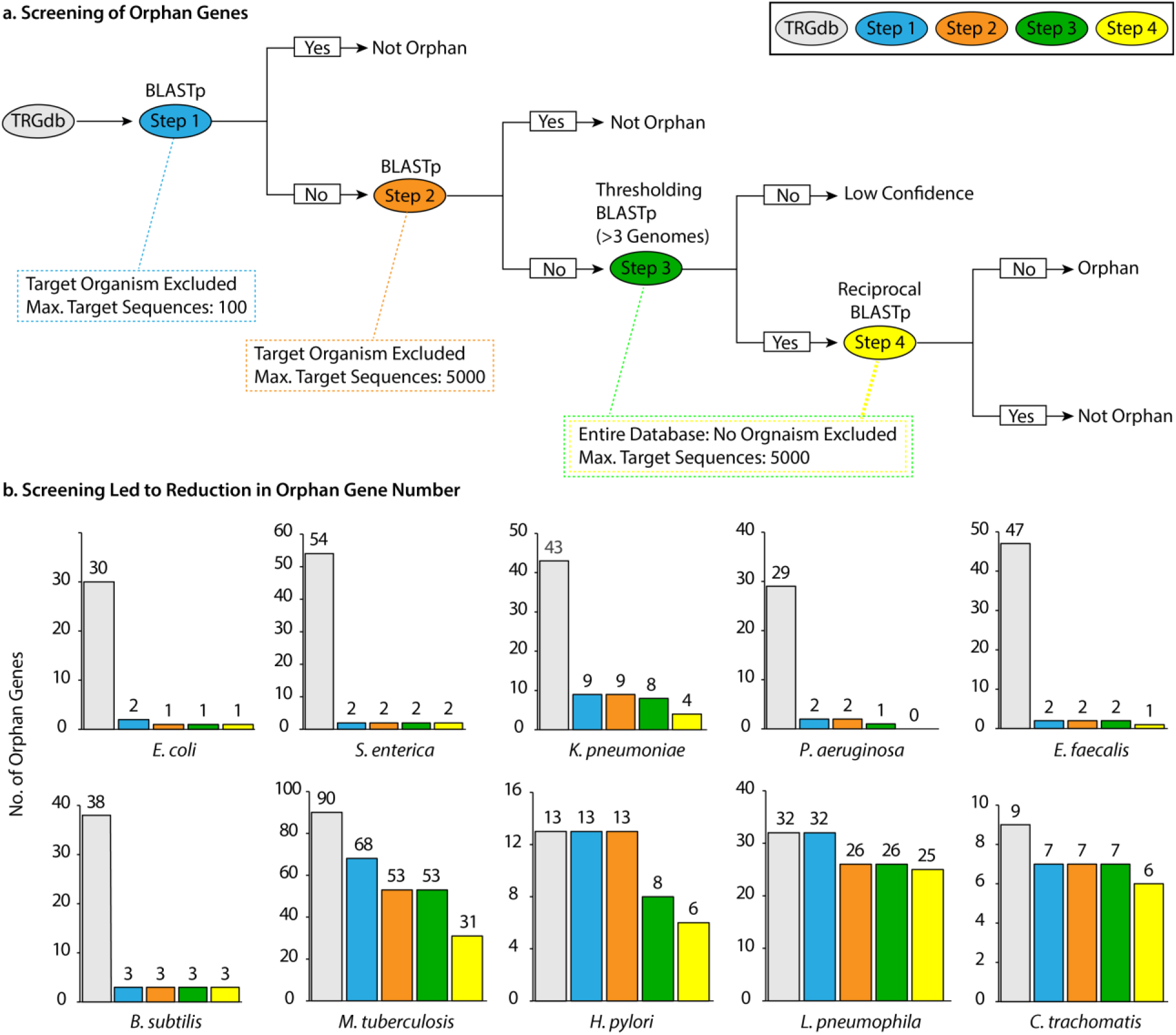
a. Schematic representation of the screening process. *Step 1*: BLASTp was performed on protein sequences obtained from TRGdb, excluding the target organism from the search. The maximum target sequences parameter was set to the default value of 100. *Step 2*: Protein sequences identified as orphan genes in *Step 1* were reanalyzed using BLASTp with an increased maximum target sequence limit of 5000. *Step 3* (**Thresholding BLASTp**): Sequences that remained orphan genes in *Step 2* were searched again, this time including the target organism. A threshold of “3 genomes” was applied, meaning that proteins detected in more than 3 genomes of the same species were considered for further analysis. *Step 4* (**Reciprocal BLASTp**): For sequences that were detected in more than 3 genomes in *Step 3*, the resulting protein hits were examined. Among these, the protein with the lowest sequence similarity to the query in *Step 3* was selected and used as the query in a reciprocal BLASTp search against the database. E value was set to 1E-3 for all four steps. **b. Reduction in orphan gene numbers across screening steps**. The screening process resulted in a progressive decrease in the number of orphan genes across 10 bacterial species: *E. coli, S. enterica, K. pneumoniae, P. aeruginosa, E. faecalis, B. subtilis, M. tuberculosis, H. pylori, L. pneumophila*, and *C. trachomatis*. A marked reduction in orphan gene numbers was observed across all species within just two years, with the most remarkable decline occurring in *Step 1*.

***Step 1:*** The initial step of BLASTp involved the exclusion of target organisms from the search, as true orphan genes should depict output as “no significant similarity found” in this search. Further, for optimizing computational efficiency and reducing analysis time, the “maximum target sequences” parameter in BLASTp was set to its default value, *i*.*e*., 100. This reduced the number of hits considered. Many genes were excluded in this step (**Fig. 2b**). Only those genes that showed no similarity at the 100-target threshold were taken to *Step 2*.

***Step 2:*** We increased the “maximum target sequences” to 5000, the maximum limit allowed by NCBI. It detected homologs for some genes that exhibited no similarity in the first round. Henceforth, the count of orphan genes declined once more in *Step 2*, inferring that the difference in result was observed when the maximum target sequences were increased to 5000 (**Fig. 2b**). We opted not to perform *Step 2* in the first place due to the intense computational demands associated with the high number of target sequences. *Step 1* facilitated quicker elimination of genes that lost their status as “orphans” by reducing processing time. The longer processing time required for *Step 2* made it more suitable as a follow-up step after narrowing down the orphan genes. The difference in results when increasing the “maximum target sequences” in BLASTp could be comprehended in how BLAST processes its search set. BLAST does not merely return the first N hits that exceed the specified E-value threshold^15^. BLAST search is divided into two phases: 1. The database search phase, where the query sequence is divided into subsequences (smaller words), and these words are then compared to target sequences in the database. 2. The alignment phase, where BLAST executes detailed alignment of these selected target sequences with the query^14^. The former step is designed to rapidly identify the most promising target sequences that are likely to produce meaningful alignments. When the “maximum target sequences” parameter is set to a lower value, the search gets restricted to a subset of the most promising sequences, optimizing speed but missing potential homologs. By increasing the value to 5000, BLAST could detect additional homologs that may not have been identified in an initial, smaller search. Though this leads to an increase in computational time, it allows for a more exhaustive comparison, thereby enhancing the sensitivity of the alignment. This emphasizes the trade-off between computational efficiency and sensitivity in database searches.

***Step 3:*** Genes that did not show any similarity in *Step 2* were further analyzed in this step. Many NCBI bacterial gene entries, including coding sequences (CDSs) and open reading frames (ORFs), originate from automated annotation pipelines that label any sufficiently long ORF as a “gene.” As a result, assembly or sequencing artifacts (e.g., misassembled contigs or repetitive regions) can produce spurious ORFs that, without RNA-seq or proteomic support, remain hypothetical and may never correspond to true, expressed proteins. To filter out these low-confidence predictions, we applied a BLASTp threshold: any ORF found in three or fewer genomes of the target species was classified as a low-confidence orphan gene. This step is crucial because bona fide orphan genes are said to evolve rapidly^16-18^ and are more likely to be conserved across multiple strains if they perform species-specific functions. In contrast, ORFs present in only a few strains often reflect horizontal gene transfer or transient mobile elements^19^ rather than stable, species-defining features. By retaining only those ORFs detected in at least four genomes, we enriched for candidates with higher functional and evolutionary relevance and substantially reduced the number of potential false orphan gene (**Fig. 2b**).

***Step 4:*** To ensure that no distant homologs slipped through our initial screens, *Step 4* employed a reciprocal BLASTp analysis^20,21^ using the ***least*** similar protein identified in *Step 3* results as the query with a E-value cutoff of 1E-3. The E-value (expected threshold) is the number of alignments with a given score that you would expect to see by chance when searching a query against the database. The threshold of 1E-3 means we kept hits whose E-value is ≤ 0.001. In other words, any match has at most a 0.1% chance of arising purely by random sequence similarity in that database. We used this cutoff because this is a common E-value cutoff^22^, and because this cutoff was used by the TRGdb in their identification of orphan genes^13^. Unlike the conventional reciprocal best-hit (RBH) approach, we deliberately used the weakest *Step 3* match to cast a wider net for remote homologs. Any gene that returned a hit with a E-value cutoff of 1E-3 to another organism in this reciprocal search was removed from our candidate list. This extra layer is vital, without it, highly divergent genes or those with low-complexity regions can masquerade as true orphans. Applying this strategy further refined our dataset, leading to an even greater reduction in orphan genes (**Fig. 2b**).

***Steps 1-4:*** Through our screening method, the four intracellular pathogens we examined (*M. tuberculosis, H. pylori, L. pneumophila*, and *C. trachomatis*) had reduced orphan genes, but this decline was smaller compared to bacteria from broader environmental niches (**Fig. 2b**). The percentage decline in orphan gene count for bacteria with broad environmental habitat ranged from 90%-100%, on the other hand, for 4 intracellular pathogen it was between 28%-66%. This observation may stem from two factors: their isolated niches, which limit opportunities for horizontal gene transfer (HGT) and genome rearrangement, and/or the smaller number of available sequenced genomes. Intracellular bacteria are known to undergo reductive evolution^23,24^, where non-essential genes are lost while highly specialized genes are retained, resulting in greater genetic stability. In contrast, bacteria inhabiting diverse environments are more prone to gene acquisitions and losses via HGT, leading to frequent changes in their orphan gene status. Although the isolation of these pathogens may contribute to the observed trend, we argue that the limited availability of sequenced genomes is likely the primary factor, as even restricted genomes showed fewer orphan genes as more genome sequencing data became available.

### General Trend: Increasing Genome Sequencing Reduces Orphan Genes

We investigated how the number of available genomes in NCBI affects the count of orphan genes across bacterial species. Bacterial species with a greater count of sequenced genomes generally exhibited fewer orphan genes. The 10 bacterial species under study were ranked in descending order based on the number of available sequenced genomes (NCBI) and divided into two groups: the top 5 possessing the highest number of sequenced genomes and the bottom 5 with the lowest (**Fig. 3, Left**). A remarkable 129-fold variation was noted in the average number of orphan genes per genome between the two groups, with the bottom five species exhibiting a notably higher count of orphan genes per genome (**Fig. 3, Right**; **Fig. S1**, individual microorganism’s orphan gene number per genome). This observation suggests that the quantity of sequenced genomes influences identification of orphan genes, probably due to enhanced annotation precision and increased genomic comparisons in well-studied species, which may result in the reclassification of formerly recognized orphan genes.

**Fig. 3.**
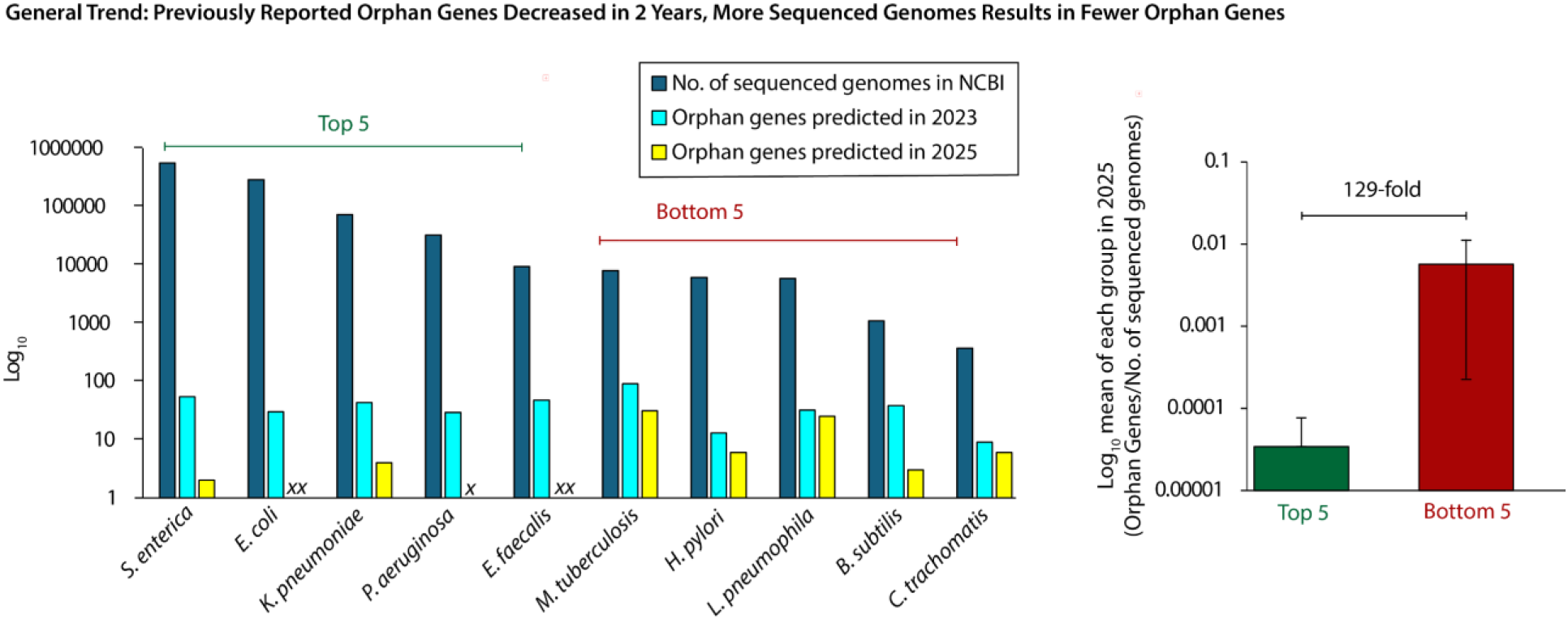
A Higher Number of Sequenced Genomes Generally Leads to a Reduction in Orphan Genes. The left panel demonstrates the number of orphan genes in 2023 and 2025, alongside the number of sequenced genomes available for each bacterial species in NCBI (**Table S1**, individual data points without log transformation). Labels: *xx*: data points corresponding to a single orphan gene (Log 1 = 0); *x*: data point corresponding to zero orphan genes (Log 0 = undefined). The 10 bacterial species studied were divided into 2 groups and arranged in descending order based on the number of genomes available (NCBI): Top 5 and Bottom 5. The right panel compares the mean number of orphan genes per genome between the Top 5 and Bottom 5. A **129-fold difference** was observed between the two groups, highlighting the impact of increased genome sequencing on the reduction of orphan genes.

Earth harbors immense microbial diversity with an estimated 10E12 microbial taxa^25^, yet scientific attention focuses disproportionately on organisms of medical or industrial relevance. The majority of microbial taxa are uninvestigated. Indeed, half of all microbiology publications center on just 10 species (*E. coli* dominates the literature), and only 26% of known bacterial species have ever been studied, leaving over 90% vastly under-investigated^26^. This uneven research landscape highlights the significant gap in scientific knowledge in microbial research and helps explain why poorly sampled species generate more apparent orphan genes.

While the number of sequenced genomes is a crucial factor, there are other factors that can contribute to the differences observed in the data. For example, *E. faecalis* and *M. tuberculosis* have similar numbers of genomes available in NCBI, yet they differ markedly in orphan genes identified in each species. This variation in orphan gene number between two species could be attributed to two factors: global prevalence and extent of genomic sampling. This is highlighted best by the World Health Organization (WHO) 2024 tuberculosis (TB) report. In this report, they showed that much of the data on *M. tuberculosis* displays a limited geographical distribution, with high prevalence in specific regions. More than 66% of worldwide TB cases originated from just eight countries: India, Indonesia, China, Philippines, Pakistan, Nigeria, Bangladesh, and the Democratic Republic of the Congo^27^. However, *M. tuberculosis* is not confined to these regions, nor to human hosts: *M. tuberculosis* has been isolated from free-ranging wildlife in South Africa^28^ and wild chimpanzee in West Africa^29^. The focus on human clinical isolates from select locales likely limits the detection of homologs in more diverse strains, inflating the number of orphan genes. In contrast, *E. faecalis*, a remarkably versatile organism, is commonly found as a commensal in the gastrointestinal tract of humans and animals^30^, and also frequently isolated from extra-enteric environments such as blood, urine, soil, water, plants, and a wide range of fermented foods (particularly in traditional dairy products)^31-34^. Furthermore, it can also be an opportunistic pathogen^30^. This widespread distribution promotes its inclusion in a broad range of studies ranging from microbiome research to clinical, environmental and food associated research. Hence, comprehensive geographical and ecological sampling is likely be responsible for strengthening genome annotation and thereby leading to fewer orphan gene identification in *E. faecalis*.

Given the immense, underexplored microbial diversity, we predict two key trends:

1. New genes classified as orphans are likely to emerge over time.
2. As more genomes are sequenced, many genes once labeled as orphans will be reassigned, perpetuating a cycle of orphan designation and subsequent reclassification. To break this cycle, we must revise both our classification criteria and our terminology. Accordingly, we propose in the next section a new framework that renames these sequences as “***Cryptogenic Gene Candidates***.”

### Revising the Terminology for Orphan Genes: A Case for “*Cryptogenic Gene Candidates*” (CGCs)

Our data demonstrated that genes once labeled as “orphan” (or taxonomically restricted) often lose their apparent uniqueness over time. We propose replacing this term with **“*Cryptogenic Gene Candidates*”** (CGCs) to more accurately reflect our current understanding. The term “Candidate” is clearer and more accurately reflects our uncertainty when no homologs are detected in nature. The term “cryptogenic,” commonly used in medicine to denote diseases of obscure or unknown origin, aptly describes genes whose origins remain uncertain. These genes could arise via *de novo* gene birth (new genes evolving from non-coding DNA) or horizontal gene transfer (the acquisition of genetic material from another organism). Although it has often been assumed that orphan genes evolve more rapidly than other genes, our new terminology acknowledges an alternative possibility: the apparent uniqueness may simply result from insufficient sequence data rather than accelerated evolution. We outline five additional reasons to retire the orphan gene and Taxonomically Restricted Genes terminology in favor of CGCs:

#### 1. Bioinformatic Artifacts

Many predicted genes might not be genuine but are artifacts of computational algorithm. The term “candidate” emphasizes the provisional nature of these findings, recognizing that further evidence would be required to confirm their uniqueness.

#### 2. Incomplete Genomic Sampling

Our understanding of genetic diversity is still tremendously inadequate. The genomes sequenced to date represent only a minute fraction of the total diversity on Earth. It is estimated that the Earth is a habitat for approximately one trillion bacterial species. Besides, fewer than one million bacterial genomes have been sequenced, the majority of which are from the same species^35^. For instance, about five years ago, it was estimated that only 2.1% of global prokaryotic taxa had been sequenced^36^. Considering predictions of up to 1 trillion microbial species and that the NCBI database currently houses approximately 330,000 prokaryotic genomes representing around 19,000 species, only roughly 2E-6% of prokaryotic genomes have been sequenced. Based on these figures, the unclear origin of many bacterial genes is primarily attributable to insufficient sequencing data, rather than the genes being truly unique.

In addition, bacterial phages, known reservoirs for gene transfer^37^, are vastly underrepresented in the genome databases. Even though phages likely outnumber bacteria by a factor of ten or more^38^. Fewer than 5000 phage genomes are cataloged in NCBI^39^. Moreover, phage genomes contain an exceptionally high proportion of novel genes with uncharacterized functions, indicating that much genetic diversity remains unexplored^38^. It has been argued that phages possess some of the greatest genetic novelty in the biological world, with as many as 80% of their encoded genes lacking similarity to known proteins and being of unknown function^38^. Thus, the challenge in tracing a gene’s origin is often more indicative of limited sequencing efforts than true genetic uniqueness.

#### 3. Limited Domain Coverage

Our study focused solely on bacteria. However, similar issues exist for archaea and eukaryotes. For example, the Genome Taxonomy Database has 584,382 bacterial genomes versus 12,477 archaeal genomes^40^, suggesting that orphan designations in archaea are likely due to under-sampling. In eukaryotes, approximately 41,000 genomes (from about 2,300 species) have been assembled^41^, yet an estimated 8.7 million eukaryotic species exist worldwide^42^. This indicates that our current data represents only about 0.026% of eukaryotic genomes in nature. Thus, the inability to trace a gene’s origins in archaea and eukaryotes is often more a reflection of limited sequencing rather than true uniqueness.

#### 4. Extinctions Lead to Limited Knowledge of Gene History

It is estimated that approximately 99.9% of all species that have ever lived on Earth are now extinct^43^. The vast majority of species went extinct before the advent of DNA sequencing. Tracing the evolutionary history of genes from extinct organisms is an extreme challenge. This gap in our data particularly affects genes that arise via *de novo* birth or horizontal gene transfer, since the donor species may be extinct. Thus, the inability to trace a gene’s origins is often more a reflection of the lack of sequenced genomes from extinct species rather than true uniqueness.

#### 5. Implications for Evolutionary Interpretations

Overestimating *de novo* gene birth rates can mislead our understanding of gene evolution and, beyond science fuel misconceptions about the origin of species. The assumption that orphan genes must have evolved extremely rapidly has even been used to argue for irreducible complexity, an argument often cited by creationists. Such conclusions rest on two flawed assumptions: first, that orphan genes will remain orphan indefinitely, and second, that any gene without a currently identifiable homolog must be unique. The limited scope of our sequencing data undermines both assumptions. Our study demonstrates that many genes previously classified as orphan genes do not retain that status indefinitely. We also showed that many genes, once considered unique due to their apparent lack of homologs in external lineages, have since been shown to share homology, indicating they are not truly lineage-specific.

Overall, the term “***Cryptogenic Gene Candidates***” **(CGCs)** more accurately encapsulates our current understanding: it highlights the uncertainty in gene origin due to incomplete genomic data rather than an inherent uniqueness. By adopting this terminology, we can foster clearer communication about what is known and what remains to be discovered in gene evolution.

### Predicting the Most Suitable CGCs for Experimental Studies

We initially retrieved 385 protein sequences from TRGdb for 10 bacterial species, and after completing all four BLASTp steps, 79 sequences remained (**Table S2**). However, four predicted proteins from *K. pneumoniae* were excluded from further analysis because the genome for this bacterial species was not fully assembled, which limited the ability to validate the presence of CGCs through this approach. As a result, genes from *K. pneumoniae* were not studied further. Of the 385 initial protein sequences, only 75 remained for further evaluation, a reduction of 81 % in potential orphan genes in only two years.

Since not all predicted genes necessarily encode functional proteins, and annotation errors can occur, it is important to assess which gene candidates would be most likely to encode truly functional proteins (**Fig. 4a**). Therefore, after completing the BLAST screening steps, we used additional computational analyses to identify the most promising CGCs for experimental validation. AlphaFold^44,45^ served as an initial filter to help identify CGCs whose protein sequences are capable of folding into stable structures. It provides an average confidence score referred to as pLDDT (predicted Local Distance Difference Test), which classifies predictions into four categories: very low (<50), low (50-70), high (70-90), and very high (>90) confidence. For our analysis, “very low” and “low” were grouped as low confidence, and “high” and “very high” as high confidence. The majority (56%) of Cryptogenic Protein Candidate sequences exhibited low structural confidence, while only 15% depicted high confidence (**Fig. 4b**). Interestingly, 29% of candidates did not produce any structural prediction (**Fig. 4b**).

**Fig. 4.**
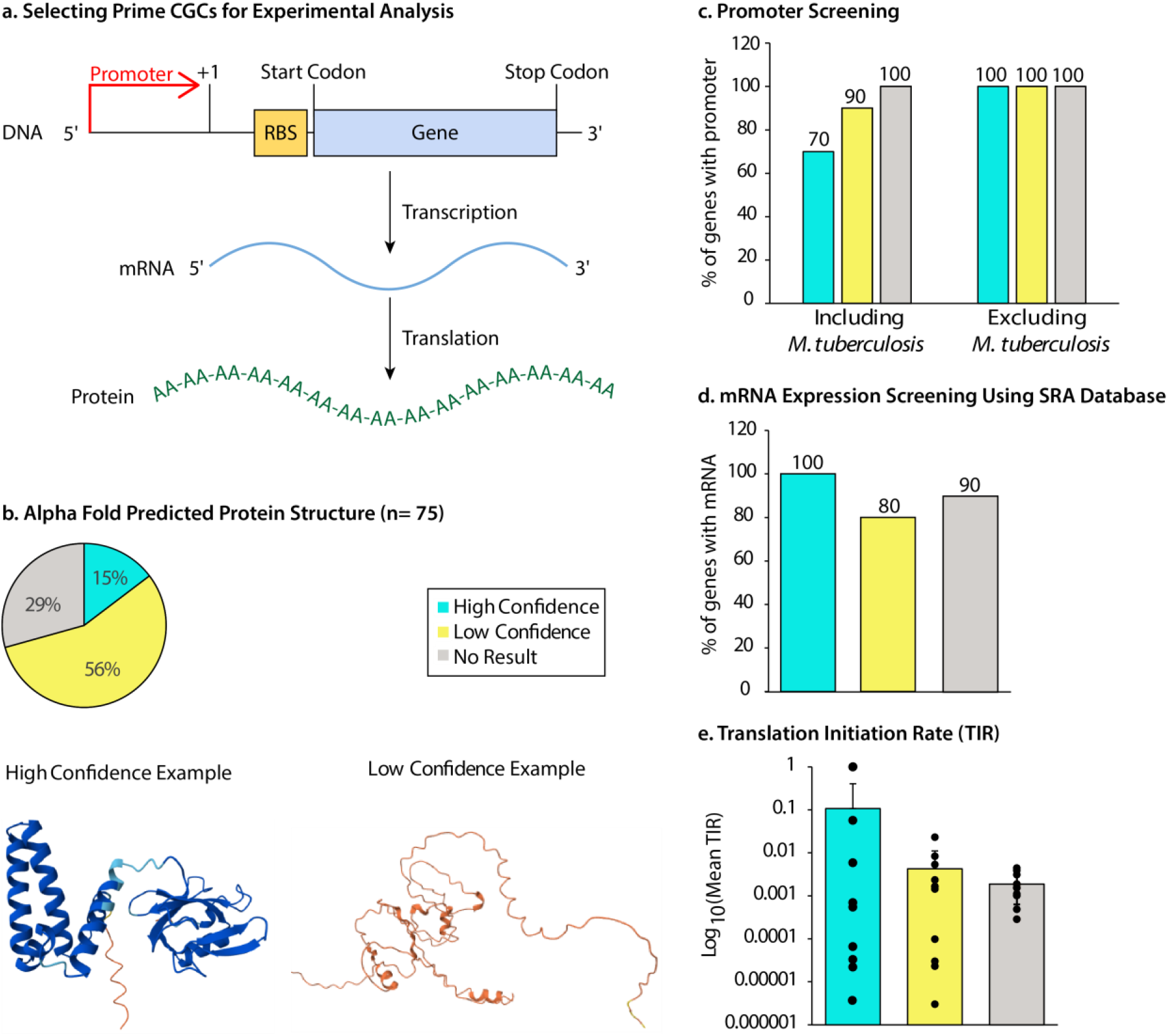
Screening of *Cryptogenic Gene Candidates* (CGCs) for experimental validation. **a**. Workflow for prioritizing CGCs based on predicted protein structure, promoter elements, mRNA expression, and ribosome-binding site (RBS) strength. **b**. AlphaFold structural predictions for 75 CGC-encoded proteins. Shown are representative models from each confidence tier: one with a high average predicted Local Distance Difference Test (pLDDT) score (90.56) and one with a low average pLDDT score (30.22). 10 proteins from each category (high confidence, low confidence, and no prediction) were selected for downstream analyses. **c**. Promoter motif search using BPROM and BioCyc. Excluding *Mycobacterium* species, every CGC across all confidence tiers contained at least one predicted promoter site. **d**. mRNA expression profiling based on RNA-seq datasets retrieved from the SRA, confirming transcript presence (but not expression level) for selected CGCs. **e**. Translation Initiation Rate (TIR) estimates. High-confidence CGCs exhibited the highest mean TIR, while the no-prediction group showed the lowest average TIR.

One reason for the high number of cryptogenic proteins with low confidence in structure could be correlated to the GC content. Codon usage is highly influenced by GC content in CGCs, which in turn plays a vital role in shaping the properties of proteins^46^. High GC genomes are likely to encode proteins that are enriched with amino acids such as alanine, proline, and glycine^46^. These amino acids tend to promote disorder, contributing to structural flexibility^47,48^. Consequently, cryptogenic proteins from high GC proportion genomes have often been indicated to be Intrinsically Disordered Proteins (IDPs). Such proteins do not possess a fixed 3D structure in solutions and exist in dynamic, flexible conformations, particularly in the absence of other proteins^49-51^. It has been revealed that low pLDDT scores might indicate the intrinsic disorder nature of proteins. Large-scale structural annotation of the human proteome using AlphaFold demonstrated that regions labeled as IDPs overlap with low-confidence predictions^52^. AlphaFold has a structure-based algorithm that is based on sequence homology to known proteins; therefore, IDPs, due to their structural plasticity, are poorly predicted by such algorithms^52-54^. Although low-confidence AlphaFold predictions do not preclude CGCs from encoding real proteins, high-confidence models enable us to prioritize those candidates most likely to succeed *in vitro* or *in vivo*.

Following AlphaFold predictions (**Fig. 4b**), 10 proteins from each category (high confidence, low confidence, and no result) were selected randomly to further investigate their likelihood of being functional genes (**Fig. 4c-e**). This analysis included checking for the presence of start and stop codons, promoter prediction, assessing mRNA expression, and evaluating translation initiation efficiency by analyzing ribosomal binding sites. All the sequences had predicted start and stop codons; however, due to the high likelihood that these codons might occur by chance, we further examined the upstream sequence of the start codon for potential promoters using BPROM.

BPROM^55^ predicted promoter regions for all candidates except those from *M. tuberculosis* (**Fig. 4c**). This software is based on the Sigma 70 promoter motif, and according to the BPROM website, it has ∼80% accuracy in predicting *E. coli* promoters. In contrast, *M. tuberculosis* does not rely on the classical Sigma 70 promoter but instead utilizes alternative sigma factors such as SigA, SigB, and SigE, which in turn recognize different promoter motifs. These motifs often diverge from the canonical -10 and -35 sequences targeted by BPROM^56-58^. Consequently, BPROM is not well-suited for identifying promoters in *M. tuberculosis*, so we also provided data where it is excluded (**Fig. 4c**). We tried predicting promoters for *M. tuberculosis* using BioCyc, but we could only predict promoters for some of *M. tuberculosis* genes. Hence, promoter screening could not be utilized as the major screening tool for experimental analysis.

To evaluate transcriptional activity of our candidate proteins, we queried the Sequence Read Archive (SRA)^59^, (a comprehensive repository of public RNA-seq datasets spanning diverse organisms and conditions) using tBLASTn to detect matching mRNA reads for each protein. While quantifying read abundance would be ideal, reconstructing RNA-seq reads across hundreds of millions of data points from different experiments and different research groups is computationally prohibitive and impractical. Rather than quantifying read abundance, we simply asked whether a given gene was expressed. Remarkably, all proteins with high-confidence structural models (100%) were supported by SRA reads, whereas 80% of low-confidence proteins and 90% of those lacking structural predictions showed evidence of transcription (**Fig. 4d**). These results demonstrate that high structural confidence is strongly predictive of mRNA expression and thus provides a powerful criterion for prioritizing candidates for experimental validation.

To assess which CGCs had potential for high translational rates (protein production rate at the ribosome), we estimated them using the Ribosomal Binding Site (RBS) Calculator^60^. On average, proteins with high structural confidence exhibited the highest predicted Translation Initiation Rates (TIRs) (**Fig. 4e**), suggesting that these proteins are more likely to be efficiently translated. These findings further support their potential for experimental validation. In summary, our integrated analysis which combines structural prediction, transcriptional evidence from the SRA database via tBLASTn, and translation initiation efficiency highlights the importance of taking into consideration of both structural confidence and expression data when prioritizing CGCs for experimental validation. Proteins with high structural confidence not only demonstrate reliable expression but are also predicted to be more efficiently translated, making them prime candidates for further investigation.

Based on our findings, AlphaFold and RBS calculator proved to be the best screening tools to identify strong experimental candidates. Here’s why: Both tools are relatively fast, reliable and can be easily automated. In contrast, the presence of start and stop codon was not informative because all gene predicting programs are based on these features. Promoter prediction proved unreliable as it can fail to predict promoters for non-model organisms like *M. tuberculosis*. In addition, the distance a promoter is from a gene is variable among organisms, making the prediction less useful. While SRA data can provide valuable expression data, it is time-consuming to analyze which makes it less practical in case of large-scale screening. Furthermore, the SRA data alone gives a binary response (yes or no in the presence of RNA), and not the quality or level of the RNA without in-depth analysis.

Through this work, we identified 10 high-confidence CGCs for experimental validation and assessed their predicted subcellular localizations. Using bioinformatic tools, we classified 1 candidate as a membrane protein, 4 as soluble proteins, 3 as secreted proteins, and 2 as signal-peptide-only fragments (**Table. S3**). The occurrence of signal-peptide-only sequences is particularly intriguing, as it suggests these peptides may represent vestigial remnants of larger proteins from which downstream domains were lost through genetic rearrangements. Further *in silico* and experimental searches for additional homologs could test this hypothesis by uncovering the missing protein segments. For example, designing universal primers to amplify homologous sequences of these peptide fragments and perform domain mapping could determine whether they integrate into larger protein products. Collectively, these findings imply that the 2 signal-peptide-only sequences may not constitute true CGCs but rather vestiges of proteins no longer functional in these bacteria.

### Implications, Future Work, and Unanswered Questions

This study relied on a database that does not incorporate the pangenome of bacterial species to identify CGCs. As a result, our analysis was limited to individual genomes rather than considering all strains within a species. Observing the trend, if the full pangenome were included, we would expect the count of CGCs to decrease further, as more homologous genes would likely be identified across strains. Due to the lack of comprehensive strain-level data, the true extent of CGCs remains uncertain. However, our results show a general trend that species with more sequenced genomes tend to have fewer CGCs.

A major challenge in orphan gene research is the frequent lack of clearly reported gene names or sequences in literature. While many studies report the percentage of orphan genes identified, they often fail to provide a list of the specific genes (by name or genes ID), even in supplementary files. In some cases, links provided turned out to be nonfunctional with the message such as “Page Not Found,” “This site can’t be reached,” or other phrases for a broken link^3,61-64^. For this study, we relied on TRGdb^13^, which offers a well-organized and searchable database, making it easier to identify genes for further analysis. Nonetheless, a significant challenge in this study was the lack of detailed gene-level information in previous research. This lack of transparency made it difficult to validate findings and perform comparative analyses.

Similarly, while analyzing bacterial promoters, we found that no single software tool was sufficient for accurate predictions across all species, requiring the use of multiple tools. This highlights the need for a more comprehensive bioinformatics tool capable of predicting promoter locations using genome sequences across a wide range of bacterial species.

Another significant challenge was analyzing gene expression data from the SRA. Although large-scale RNA-seq datasets are available, identifying accession numbers that match the exact strain used in our study proved difficult. Due to feasibility constraints, we limited our analysis to a small subset of accessions at a time. This highlights the need for a dedicated software platform that categorizes RNA-seq datasets at the species and strain level, making them more accessible for future research. Additionally, our analysis focused solely on determining whether the CGC sequences were transcriptionally active without taking into account expression levels, experimental conditions, or potential regulatory influences.

A further issue arises from the way BLAST/NCBI handles result ordering when excluding a specific organism. For example, in our *E. coli* screening pipeline, *Step 1* requested the top 100 hits while excluding *E. coli*, yielding no hits from any other species. In contrast, *Step 2* used the top 5000 hits with *E. coli* excluded and did return matches from other organisms. This discrepancy occurs because BLAST first compiles all potential hits (the top were *E. coli* sequences), and only afterward removes those belonging to the excluded taxon. In *Step 1*, the first 100 results were entirely *E. coli*, so once those were filtered out, nothing remained. By expanding the search to 5000 hits in *Step 2*, BLAST was able to gather enough non–*E. coli* matches to populate the filtered set.

While this workaround served our immediate needs, it has inherent limitations. As genomic databases continue to grow at an exponential pace, it will soon be common to see more than 5000 high-scoring *E. coli* sequences for a given gene. At that point, even requesting 5000 hits will still capture only *E. coli*, and any more distantly related homologues, though present, will be excluded because they never enter the initial pool of top matches. In principle, one could raise the cutoff to 50,000 or beyond to ensure coverage of distant homologs, but NCBI currently caps BLAST results at 5000 entries. To make CGC identification and other evolutionary analyses more robust and accessible to the broader research community, BLAST/NCBI will need to either raise this 5000-hit ceiling or implement alternative filtering strategies that avoid this queueing limitation. A change needs to be implemented soon because the NCBI genomic repository is growing at an extraordinary pace.

In selecting CGCs for experimental validation, we prioritized proteins with high-confidence structural predictions from AlphaFold, particularly because these candidates also exhibited strong transcriptional evidence—such as the presence of mRNA, promoters, and higher average Translation Initiation Rates. These indicators support the likelihood that these genes are expressed and translated into functional proteins, making them suitable starting points for downstream characterization. However, a substantial proportion of the CGCs that yielded low-confidence or no structural predictions also showed clear evidence of transcriptional activity, including mRNA, promoters and TIRs. This suggests that these genes may still be expressed and potentially encode proteins, despite their limited or uncertain structural predictability. It is important to emphasize that the absence of a confident AlphaFold prediction does not imply that a gene is non-coding. Rather, it may reflect the novelty or divergence of the sequence, which falls outside the model’s trained structural space. Therefore, while our current strategy focuses on high-confidence AlphaFold predictions for initial experimental validation, we do not exclude low-confidence or no-result candidates from being legitimate protein-coding genes.

These challenges align with broader issues in microbial genomics. A recent study determined that among the studied bacterial species, a disproportionate amount of research focuses on just 10 species, with more than 90% of bacteriology articles examining less than 1% of all species^26^. This limited representation affects our understanding of CGCs. In other words, as we sequence more genomes, we are likely to uncover additional orphan genes as genomic datasets from poorly studied or underrepresented species often do not contain sufficient annotation to recognize homologous genes, leading to an overestimation of orphan genes in earlier studies; however, with continued sequencing, genes once considered unique to a specific lineage may no longer be viewed as distinct.

We currently lack a comprehensive study confirming that some CGCs represent genuine, protein-encoding genes. While our analysis has highlighted several promising candidates, it is equally plausible that some (of the good or not so good candidates) may not encode functional proteins. Moving forward, it is crucial to experimentally investigate the roles of the proteins these genes encode, whether through gene knockout, overexpression studies, or innovative alternative methods. Such functional analyses will offer deeper insights into the biological significance of CGCs and their potential contributions to bacterial adaptation and evolution.

## Conclusion

In this study, we reassessed orphan genes using a BLAST-based approach to determine their presence across bacterial genomes. Our findings indicate that the number of CGCs is influenced by the availability of sequenced genomes, with a general trend of decreasing CGCs counts as more genomes are included. This suggests that many genes initially classified as orphans may simply lack annotated homologs due to incomplete genomic data rather than being truly unique to a species. However, the absence of enough genome sequences (from bacteria to viruses) mean that our estimates may still be inflated. Beyond genomic presence, we encountered several challenges in studying CGCs, including the lack of gene-level data in previous studies, the limitations of available promoter prediction tools, and the difficulty in linking RNA-seq datasets to specific bacterial strains. These challenges highlight the need for improved bioinformatics tools to facilitate more precise gene annotation, promoter identification, and transcriptomic analysis at the strain level. Future research should focus on integrating pangenome analyses to refine orphan gene classification and reduce overestimation. Furthermore, functional validation through gene knockout and overexpression studies will be essential for understanding the biological roles of these genes. By expanding sequencing efforts and improving computational tools, we can gain deeper insights into the evolutionary significance of CGCs and their potential contributions to bacterial adaptation and innovation.

## Methods

### Data Retrieval and Selection

To investigate orphan genes in bacterial species, we obtained protein sequences from the Taxonomically Restricted Genes Database^13^ (TRGdb) which contains taxonomically restricted genes from 80,789 bacterial species at the genus and species levels. We selected 10 bacterial species for this study and retrieved their corresponding orphan protein sequences (**Table S4**). The bacterial species and their respective genome accession numbers are listed in **Table 1**.

**Table 1.** A list of bacterial species and genome accession numbers used in this study.

| Sr. No. | Bacteria Species | Genome Accession No. |
| --- | --- | --- |
| 1 | *Bacillus subtilis* | GCF_000009045 |
| 2 | *Chlamydia trachomatis* | GCF_000012125 |
| 3 | *Enterococcus faecalis* | GCF_000392875 |
| 4 | *Escherichia coli* | GCF_003697165 |
| 5 | *Helicobacter pylori* | GCF_001653475 |
| 6 | *Klebsiella pneumoniae* | GCF_000742135 |
| 7 | *Legionella pneumophila* | GCF_000008485 |
| 8 | *Mycobacterium tuberculosis* | GCF_000195955 |
| 9 | *Pseudomonas aeruginosa* | GCF_001457615 |
| 10 | *Salmonella enterica* | GCF_000006945 |

### Orphan Genes Screening Using BLASTp

The screening of orphan gene (CGCs) was conducted using a four-step BLASTp^14^ approach, with an expected threshold of 1E-3 for all four steps, consistent with the TRGdb classification criteria for orphan genes. In *Step 1*, BLASTp was performed on the protein sequences of each bacterial species from TRGdb, excluding the target organism, and the maximum target sequences set to 100 (default value). A protein would be classified as an orphan if it showed no significant similarity to any sequence in the database. Henceforth, protein sequences that exhibited no significant similarity in *Step 1* were subjected to *Step 2*, where BLASTp was performed again. This time maximum target sequences were increased to 5000. In *Step 3*, termed thresholding BLASTp, sequences that showed no significant similarity in *Step 2* were reanalyzed, and the target organism was reintroduced while maintaining a maximum target sequences value 5000. Genes found in three or less genomes within the target organism’s species were classified as low-confidence orphan genes (excluded from further analysis), while those in more than three genomes were classified as high-confidence orphan genes. In *Step 4*, a reciprocal BLASTp search was conducted to identify potential hidden homologs for high-confidence orphan genes. The protein sequences with the lowest similarity scores from *Step 3* results were used as queries against the complete BLASTp database. It also involved the inclusion of the target organism, with a maximum target sequence value set to 5000.

### Structural Analysis Using AlphaFold 3

AlphaFold^45^ was then used to predict the structures of 75 proteins obtained in the end (**Table S5**). The process begins with multiple sequence alignment (MSA) which is based on homology, followed by the prediction of protein structures. The confidence of each prediction is assessed using the predicted Local Distance Difference Test (pLDDT) score. The results were categorized into three groups: high-confidence (including very high and high pLDDT scores), low-confidence (low and very low pLDDT scores), and no result (for sequences without structural predictions).

### Screening of Orphan Gene Presence Using Custom tBLASTn

After AlphaFold was used to predict the structures of the 75 proteins, we wanted to confirm their actual presence in the respective genomes as mentioned in TRGdb before proceeding with further computational analyses. To do this, a custom tBLASTn analysis was performed using Geneious Prime software. This step ensured that the genes were correctly annotated in the genomes of the selected bacterial species, thereby ruling out the possibility of potential annotation errors. The genome sequences of each bacterial species were obtained in “.gbff” and “.fasta” formats from NCBI and imported into Geneious bioinformatics software. A Custom tBLASTn search was conducted on ten proteins from each confidence category (high-confidence, low-confidence, and no result). Proteins that did not show 100% identity or 100% query coverage were excluded from further analysis. These proteins were replaced by other candidates from the same category that met the 100% identity and query coverage criteria. This validation confirmed the presence of orphan genes in the genomes, ensuring that no annotation errors were present before proceeding with further computational studies. This also provided the location of an orphan gene within the genome which enabled us to retrieve nucleotide sequence corresponding to protein sequence (**Table S6**).

### Functional Analysis of *Cryptogenic Gene Candidates*

To predict whether the CGCs encode functional proteins or are mere annotation artifacts, we proceeded with additional analyses on 10 randomly selected candidates from each confidence category (high-confidence, low-confidence, and no result) (**Table S6**). For each candidate, we assessed the presence of key transcriptional and translational features, including predicted promoter regions, mRNA transcripts, and Ribosome Binding Sites (RBS). This multi-layered approach allowed us to better understand the potential expression and coding capacity of CGCs across different structural confidence levels.

### Promoter Region Detection Using BPROM

The first step in assessing functionality was to check whether the genes were likely to be transcribed. We used BPROM^55^, a tool that identifies potential promoter regions in bacterial genomes. Using BPROM, we analyzed a 100 bp upstream sequence from the start codon for each gene (**Table S6**) and predicted the presence of a promoter. A confirmed promoter region suggests that the gene has the potential to be transcribed into mRNA. If no promoter was detected, we proceeded to the next step to investigate the possibility that the gene could be part of an operon.

### Operon Structure Analysis Using BioCyc

If no promoter was detected, the next step was to explore whether the CGCs might be part of an operon. The presence of an operon is a common feature in prokaryotic genomes where groups of genes are co-transcribed as a single mRNA molecule. We used BioCyc^65^ to examine whether the gene could be part of an operon. It is a curated database that contains detailed information on metabolic pathways and operon structures. A gene within an operon may still be transcribed even without a clear promoter region.

### mRNA Expression Validation Using SRA Data

To analyze whether the CGCs were actively expressed and not just annotated without functional support, we checked for mRNA expression. This was done using the NCBI Sequence Read Archive (SRA), a database that contains RNA sequencing datasets from a variety of bacterial strains. For each selected gene, we performed tBLASTn searches against RNA-seq data corresponding to the strain in which the CGCs was identified. We used the SRA accession numbers (**Table S6**) to search for RNA-seq data, ensuring that we used datasets from the correct strain. If no hits were obtained after testing three different SRA accessions, the gene was considered to have no detectable mRNA expression. This implies that it might not be actively transcribed or functional.

### Ribosome Binding Site (RBS) Analysis for Translation Potential

To evaluate whether the CGCs had the potential for translation, we analyzed their Ribosome Binding Sites (RBS). The RBS is a critical component for initiating translation, as it guides the ribosome to the start codon. To estimate the translation initiation potential, we used the RBS Calculator from Salis Lab^60^. We input the nucleotide sequence of the protein-coding region of the gene, along with 50 bp upstream, into the RBS Calculator. This tool estimates the strength of the RBS based on its thermodynamic properties. The calculator provides a total free energy value: lower free energy values indicate stronger ribosome binding and higher translation potential, while higher free energy values suggest weaker binding and reduced translation efficiency. This step helped us assess whether the CGCs were likely to be translated into functional proteins.

Through this detailed analysis that included promoter region detection, operon structure analysis, mRNA expression validation, and RBS evaluation, we were able to determine whether the CGCs were likely to encode functional proteins or if they were likely to be non-functional or annotation artifacts. This rigorous validation process ensured that the genes selected for further analysis had the potential to be biologically relevant and functional.

### Protein Property Prediction and Membrane Localization Analysis

We performed *in silico* analysis employing bioinformatic tools to predcit the solubility and association of proteins with membrane for 10 high confidence CGCs (**Table S3**). This step was desired to identify proteins which would favour downstream processing such as expression and purification. Protein sequences corresponding to desired CGCs were analyzed using ProtParam^66^, ProtScale^66^ and DeepTMHMM^67^. ProtParam predicts certain physiochemical properties such as theoretical pI, instability index, grand average of hydropathicity (GRAVY), etc. GRAVY values estimate overall hydrophobic or hydrophilic nature of a protein: positive values stipulate hydrophobicity and negative values indicate hydrophilic nature. This provided us the information related to solubility in aqueous environments. Then, ProtScale tool from the ExPASy was utilised to assess membrane association of proteins. A window size of 19 was applied, which is commonly used to detect transmembrane helices^66^. The Kyte & Doolittle hydrophobicity scale was selected, and the output predicts the hydrophibicity profile, where peaks exceeding a value of 1.6 suggested the presence of potential transmembrane segments. To predict the presence of transmembrane region, we applied DeepTMHMM tool. It classifies each region of protein as cytoplasmic, extracellular, signal peptide, or membrane embedded.

## Authors’ Contributions

SKM conducted all analyses, prepared the manuscript, and created the figures. XYB developed scripts to extract sequences of the analyzed bacterial species from the TRGdb.fasta file, which contains protein sequences from 80,789 bacterial species, totaling 247,617,414 sequences. SXG contributed to the conceptualization of using SRA and AlphaFold to assess whether the identified sequences express mRNA and form stable structures, respectively, and guided the analysis. NCB formalized the research concept and oversaw the project. All authors contributed to manuscript discussions and revisions.

## Data Availability

All the data that supports the findings of this study are included in this published article (and its supplementary information files). Also, any additional data is available from the corresponding author upon request.

