## Supplementary material for "From Orphan Genes to Cryptogenic Gene Candidates: Reassessing Uniqueness": fig._s1_v2.docx

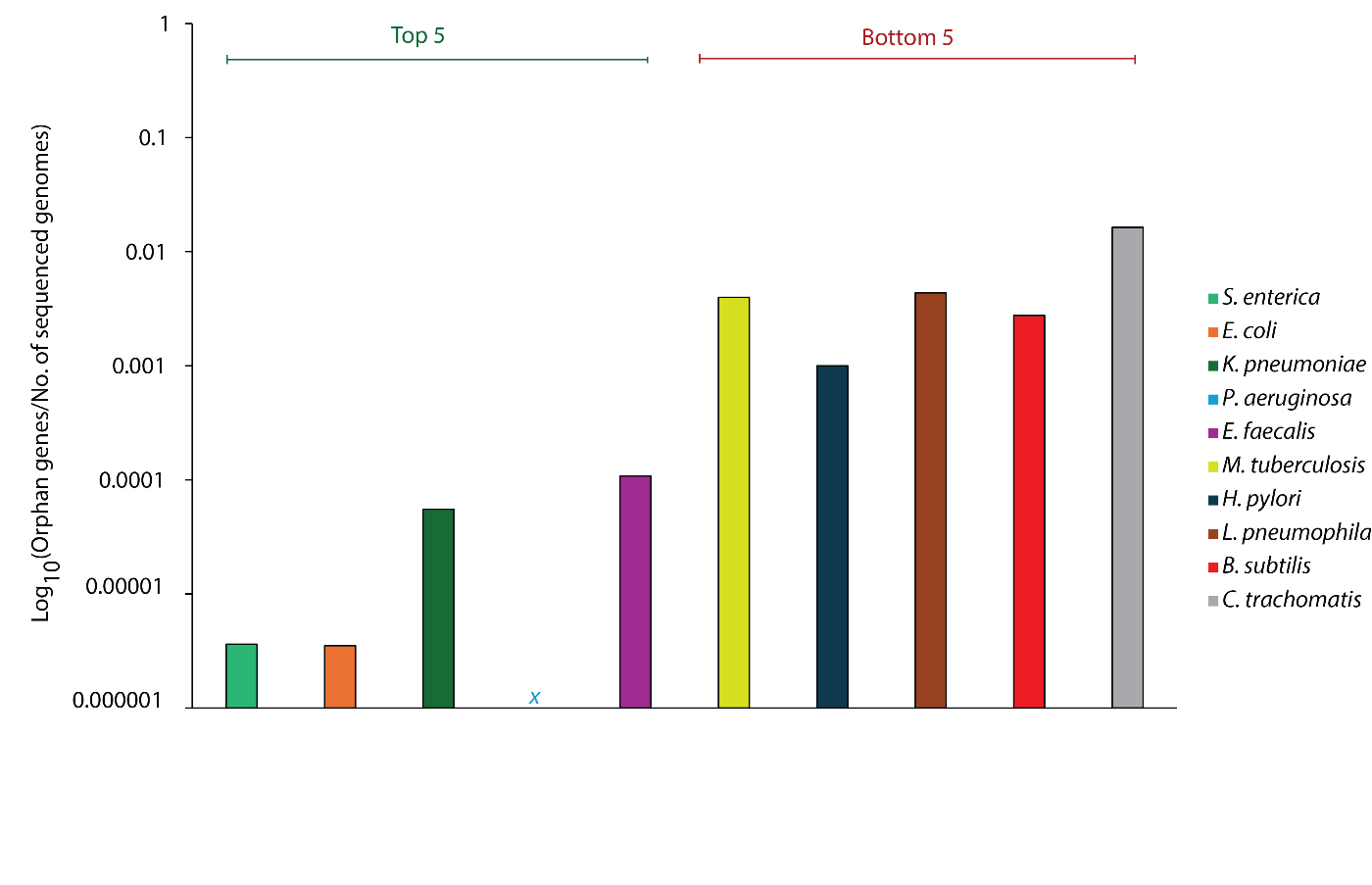

**Fig. S1.** **Ratio of orphan genes to the number of sequenced genomes for each microorganism.** The plot displays the top five and bottom five groups based on this ratio. Label: *x*: data point corresponding to zero orphan genes (Log 0 = undefined).
